# Major depressive disorder recurrence and medication status shape brain network topology

**DOI:** 10.64898/2026.09.18.752576

**Authors:** Javier F. Castilla-Jiménez, Juan Carlos Díaz-Patiño, Stefanie Enriquez-Geppert, Marie-José van Tol, Fernando A. Barrios, the DIRECT Consortium, Sarael Alcauter

## Abstract

**Introduction:** Major depressive disorder (MDD) is a highly prevalent and disabling psychiatric disorder. Human neuroimaging studies increasingly frame its neurobiological substrate in terms of alterations of large-scale brain network organization. Resting-state fMRI findings broadly align with this view, yet remaining highly heterogeneous, reflecting both clinical and analytic variability. Predominant approaches largely characterize functional organization in terms of pairwise relationships between regions, which may limit the ability to capture more global features of brain organization associated with depression and its course. Here we use novel methods from Topological Data Analysis to provide new perspectives on these brain-behavior associations

**Objectives:** The main objective of this study was to determine whether whole-brain topological descriptors of functional connectivity capture alterations associated with MDD. In particular, we examined whether these features vary along clinically relevant dimensions of heterogeneity, focusing on illness course (single-episode vs. recurrent MDD) and current antidepressant medication status. To this end, we analyzed the area under the curve (AUC) of the zeroth and first Betti numbers (B₀ and B₁), which index global network integration and higher-order cycle structure across scales, respectively.

**Methods:** Neuroimaging (resting state fMRI) and phenotypic data were obtained from 1,490 participants (776 individuals with MDD and 714 never-depressed controls) included in the REST-meta-MDD project of the DIRECT consortium. Functional connectivity matrices were computed using Pearson correlations between regions defined by the Power-264 atlas and harmonized across sites using CovBat. Topological Data Analysis was applied to characterize whole-brain network organization across connectivity thresholds using B₀ and B₁ curves and their areas under the curve (AUC). B₁ associated cycle counts and their coarse anatomical configurations were further examined. Group differences and effects of illness course and medication status were assessed using multiple linear and generalized linear regression models.

**Results:** Compared with controls, individuals with MDD showed significantly higher B₀ AUC, indicating altered global integration profiles across connectivity thresholds. B₀ AUC did not differ by illness course, illness duration, or symptom severity. In contrast, B₁ AUC showed a graded increase across illness courses, with the highest values observed in recurrent depression. This effect was driven by an increased number of one-dimensional cycles rather than greater cycle persistence. Medication status further modulated B₁ alterations, with elevated values observed in unmedicated single-episode and medicated recurrent MDD. Anatomical decomposition revealed that higher-order alterations were primarily driven by inter-network configurations spanning multiple large-scale functional systems.

**Discussion:** These findings indicate that MDD is associated with altered multiscale functional organization, combining less compact global integration with an increased prevalence of cross-network hole-defining cycles, with these features varying systematically with illness course and current medication, suggesting that topology-informed measures capture clinically relevant variability in brain organization in depression.

## Introduction

Major depressive disorder (MDD) is a highly prevalent and disabling psychiatric disorder (American Psychiatric Association, 2013; World Health Organization, 2017), with a global lifetime prevalence of approximately 10% (Lim et al., 2018). Despite the availability of evidence-based treatments, many individuals with MDD do not achieve sustained remission (McIntyre et al., 2014; Papakostas and Fava, 2008). Recurrent depressive disorder (MDD-R), is characterized by repeated episodes of depression (World Health Organization, 2024) and is associated with a more severe and disabling clinical course than single-episode depressive disorder (MDD-SE) (Ahern et al., 2025; Crowe et al., 2020; Roca et al., 2011; Semkovska et al., 2019; Varghese et al., 2022). Nonetheless, the pathophysiological mechanisms underlying MDD and its recurrence remain poorly understood (Pilmeyer et al., 2022).

The etiology of MDD is multifactorial and is commonly framed within a biopsychosocial model, encompassing interacting biological, psychological, and social influences (Otte et al., 2016; Remes et al., 2021). This model reflects the current consensus that multiple interacting factors contribute to depressive pathology, without a single unified mechanism (Cui et al., 2024; Malhi and Mann, 2018; Otte et al., 2016). These factors, however, do not operate uniformly across the course of illness. Evidence suggests that mechanisms associated with first onset and those governing recurrence only partially overlap; while sex is strongly linked to first-episode onset, it appears less informative for recurrence, where risk profiles are largely shared across sexes (Van Loo et al., 2018). Conversely, factors more closely tied to recurrence, such as prior episodes and residual symptoms, differ from those implicated in initial onset and are further shaped by the treatment trajectory (Buckman et al., 2018).

The brain provides a convergent level at which these diverse influences may be expressed and studied, without assuming that any single brain abnormality constitutes a unitary cause of MDD. Consistent with this view, a growing body of research suggests that core depressive symptoms are associated with widespread abnormalities across functionally related brain systems, with effects that extend beyond isolated regional responses (Javaheripour et al., 2021; Kaiser et al., 2015; Yang et al., 2021). This line of work has motivated the neuroscientific conceptualization of MDD as a network-based brain disorder (Duran and Miller, 2020; Li et al., 2018, 2023).

Resting-state functional magnetic resonance imaging (rs-fMRI), enables the study of spontaneous brain activity, and has become a central tool for probing large-scale network alterations in MDD (Castellanos et al., 2013; Zhang et al., 2025). Early rs-fMRI studies, together with subsequent meta-analyses, suggest that depressive symptoms are linked to distributed abnormalities in intrinsic connectivity, particularly within the default-mode, cognitive control and salience networks, but also highlight substantial variability in the spatial patterns and directions of these effects across studies (Kaiser et al., 2015; Mulders et al., 2015; Zhang et al., 2025). Several factors contribute to the heterogeneity observed across rs-fMRI studies of MDD. At the clinical level, MDD is defined by a broad constellation of symptoms and course characteristics, such that individuals may meet diagnostic criteria while sharing only a limited subset of features (Buch and Liston, 2021). This variability is reflected in clinical specifiers (e.g. anxious distress, atypical symptom, or psychotic features) that modulate symptom expression and prognosis across individuals (Lamers et al., 2016; Nelson et al., 2018; Zimmerman et al., 2019).

Beyond symptom-level specifiers, the course of illness delineates broader trajectories of disease progression: in particular, the distinction between single-episode and recurrent forms of MDD has been consistently associated with differences in symptom severity, cognitive burden, functional impairment, and treatment resistance, while acknowledging substantial heterogeneity within each course-defined group (Ahern et al., 2025; Crowe et al., 2020; Roca et al., 2011; Semkovska et al., 2019; Varghese et al., 2022). While modest sample sizes are repeatedly cited as major barriers to consistency across rs-fMRI studies (Li et al., 2022; Turner et al., 2018), increasing sample size alone does not resolve the heterogeneity inherent to MDD. Large-scale initiatives designed to increase statistical power have reported distributed and sometimes partially overlapping alterations, with no single connectivity pattern that generalizes across all patients. For example, the REST-meta-MDD consortium, reported reduced default-mode connectivity in recurrent MDD, along with alterations in multiple systems (Chen et al., 2022; Yan et al., 2019), whereas other multisite study found concurrent increases in default-mode connectivity and decreases in sensorimotor and visual systems in MDD (Zhang et al., 2024a). In contrast, the PsyMRI consortium reported only subtle, spatially distributed group differences without a robust canonical pattern (Javaheripour et al., 2021).

Beyond clinical and sample-size considerations, heterogeneity in rs-fMRI findings is further amplified by differences in analytic choices (Botvinik-Nezer et al., 2020). Recent work has shown that analytic decisions (including preprocessing, parcellation and connectivity definition), can meaningfully alter functional-connectivity estimates, even within the same dataset (Alakörkkö et al., 2017; Bielczyk et al., 2018; Botvinik-Nezer et al., 2020; Dimitriadis et al., 2017; Kopal et al., 2020; Luppi et al., 2024; Parkes et al., 2018; Pavlovich et al., 2025), a pattern echoed by recent reviews and meta-analysis of rs-fMRI studies in depression reporting substantial variability across otherwise comparable studies (Li et al., 2022; Pilmeyer et al., 2022; Sundermann et al., 2014; Zhang et al., 2025). Within this landscape, network neuroscience largely relies on approaches (e.g. graph theory) that summarize functional organization in terms of pairwise relationships between brain regions (Centeno et al., 2022). While these methods provide important insights, they limit the ability to capture how groups of regions jointly organize and to further describe the characteristics of such larger structures (Sizemore et al., 2019).

Topological Data Analysis (TDA) and in particular persistent homology, provides a complementary framework by characterizing network organization in terms of global structural features, such as the integration into connected components or the presence of higher-order cycles (Centeno et al., 2022; Giusti et al., 2015; Salch et al., 2021; Sizemore et al., 2019). Rather than focusing on individual connections, persistent homology quantifies how these global features emerge and persist, as the overall structure of the network is examined across a continuum of sparsity thresholds, providing a multiscale summary of how frequently such features occur and how stable they are within the system. This approach offers a principled way to separate robust system-level structure from pipeline-dependent variability. Recent work shows that topological descriptors are robust to variability of acquisition (Kumar et al., 2023) and brain parcellation (Gracia-Tabuenca et al., 2020), while being sensitive to developmental (Gracia-Tabuenca et al., 2023) and clinically relevant differences in neuropsychiatric conditions such as ADHD and schizophrenia (Centeno et al., 2022; Gracia-Tabuenca et al., 2020; Salch et al., 2021).

To operationalize this approach, we leverage the REST-meta-MDD consortium database, to calculate Betti curves, compact descriptors of how topological features behave across different levels of connectivity, and base our analyses on summary measures extracted from these curves. We test whether individuals with MDD show alterations in topological descriptors of whole-brain functional organization. A central aim of the present study is to determine whether this topological characterization captures clinically relevant heterogeneity in depression, particularly illness course. Specifically, we assess differences between recurrent MDD and single-episode MDD, as well as the influence of current medication status. Based on prior evidence, we hypothesize that recurrent MDD will be associated with more pronounced alterations in global topological organization, reflecting differences in the integration and higher-order structure of large-scale networks. By addressing these questions, the present work aims to determine whether topology-informed descriptions of brain organization can add a complementary layer of understanding to individual variability in the course of depression.

## Methods

### Sample composition

Resting-state fMRI and phenotypic data were used from the REST-meta-MDD project (Yan et al., 2019), part of the DIRECT (Depression Imaging REsearch ConsorTium) initiative. The full dataset comprises 1,300 individuals diagnosed with MDD and 1,128 age-matched never-depressed controls, collected across 25 sites in China. Psychiatric diagnoses were established using ICD-10 or DSM-IV criteria. Phenotypic data included sex, age, and education years for most participants, and, when available, also medication status at scan time, illness duration, illness course (single-episode vs. recurrent), and symptom severity (assessed via either the 17- or 24-item Hamilton Depression Rating Scale, HAMD). Ethical approval for the collection of the original REST-meta-MDD dataset was granted by the Institutional Review Board (IRB) of the Institute of Psychology, Chinese Academy of Sciences (approval No. H19045), and all participants provided written informed consent prior to participation. The present study constitutes a secondary analysis of de-identified data obtained through the DIRECT consortium in accordance with the consortium’s data-sharing procedures.

The inclusion and exclusion criteria (see supplementary Fig. S1) described by Yan et al. (2019) were used, excluding participants younger than 18 or older than 65 years, individuals with missing demographic or imaging data, those with poor spatial normalization, and those with excessive head motion (mean framewise displacement > 0.2 mm). In addition, data labeled as belonging to site 4 of acquisition was excluded owing to overlap with data from site 14, as indicated in the dataset documentation. After this filtering, data of 1,601 participants remained (830 MDD and 771 NDC), and from these, 1,490 individuals (776 MDD and 714 NDC; 15 sites) had complete data for the Power’s 264 functional atlas (Power et al., 2011). This final full sample was used for all subsequent analyses. Three analytic working samples (Table 1), with comparable demographics (Fig. 1 and supplementary Fig. S2) were defined according to the availability of clinical metadata relevant to each research question:

1. For comparisons by diagnosis (MDD vs. Control), the final full sample (**n = 1,490**) was used.
2. For analyses by illness course (single-episode MDD [MDD-SE] vs. recurrent MDD [MDD-R]), a subsample was defined by discarding the subset of MDD participants with no available recurrence data (working sample **n = 1,296**, including all controls).
3. For analyses of current (antidepressant) medication status, a subsample was defined by discarding the subset of MDD participants with both unavailable recurrence data and unavailable current medication status (working sample **n = 1,166**, including all controls). MDD participants in this subsample were classified into four subgroups: medicated MDD-R (mMDD-R), unmedicated MDD-R (uMDD-R), medicated MDD-SE (mMDD-SE), and unmedicated MDD-SE (uMDD-SE) (see Table 1).

**Fig. 1.**
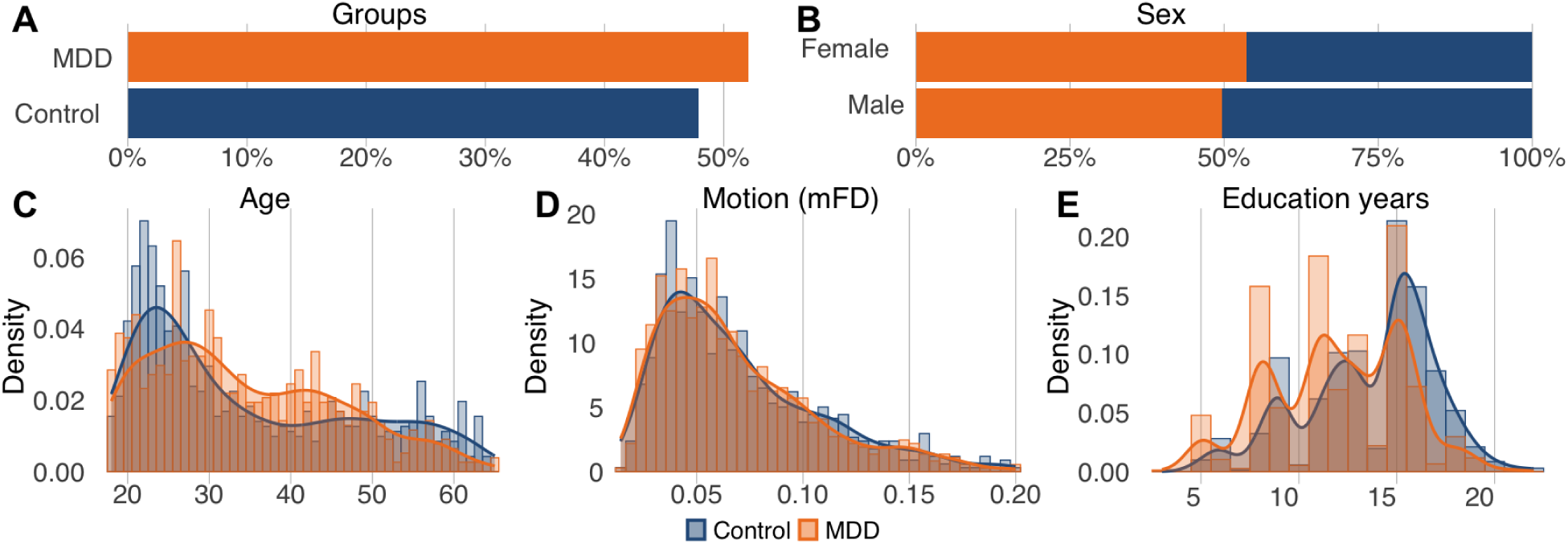
Demographic characteristics of the final full sample. (**A**) Groups proportions by diagnosis, (**B**) Sex proportions by diagnosis, (**C**) Age density distribution by diagnosis, (**D**) mean head motion (mean framewise displacement, mFD) by diagnosis, and (**E**) education years by diagnosis, are shown for the final sample as used in diagnosis-level analyses (n = 1,490; 776 MDD and 714 Controls). Demographic variables were comparable across all analytic subsamples and were included as covariates in all the corresponding statistical models. Graphs of demographics characteristics at the subsample level are provided in Supplementary Fig. S2.

**Table 1.**
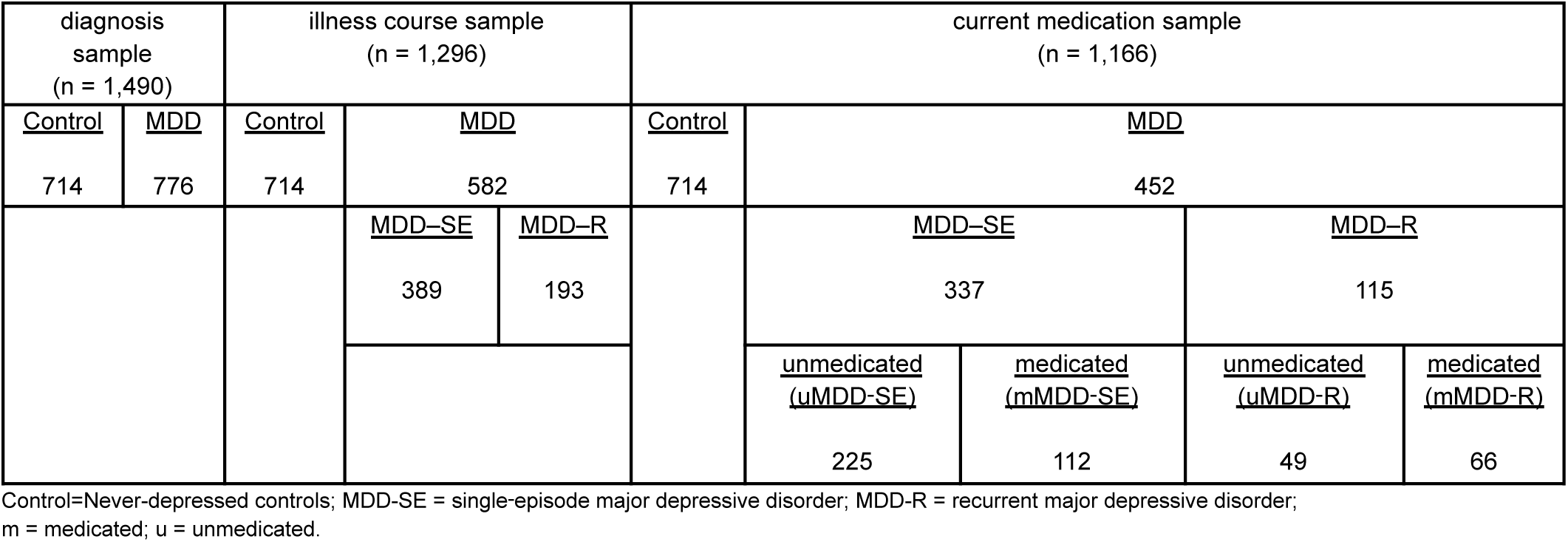
Working sample composition across diagnostic, illness course, and current medication analyses.

For secondary analyses assessing associations between topological indices and either illness duration (in weeks) or clinical severity (HAMD scores) within the recurrence-defined subsample, only MDD participants with available data on either illness duration or HAMD scores, were retained. These variables were only available for a limited subset of MDD participants, and the corresponding sample sizes are reported alongside the results.

### fMRI preprocessing

All scans were locally preprocessed at each site using an identical protocol, as described by Yan et al., 2019. Preprocessing steps included slice-timing correction, realignment, nuisance regression, spatial normalization, and signal filtering. The consortium provided final resting-state fMRI indices, defined as the time series of the average signal per region, which were generated either with or without applying global signal regression. To ensure robustness, all analyses were performed using the dataset with global signal regression, as it has been shown to be an effective approach to reduce nuisance confounds from fMRI data (Xifra-Porxas et al., 2021).

### Functional connectomes

Power’s 264 (P264) functional brain atlas (Power et al., 2011) was used by the REST-meta-MDD consortium during preprocessing to segment the brain into 264 regions of interest (ROIs), from which average regional time series were extracted and provided as part of the released dataset (Yan et al., 2019). Using these ROI time series, static functional connectivity matrices were computed for each subject by calculating Pearson correlation coefficients between all pairs of ROIs. By this process a whole-brain connectome was computed for each subject. All functional connectome calculations were performed using the R statistical computing environment (R Core Team, 2024).

### Harmonization

Due to the multi-site nature of the dataset, fMRI scans were acquired using different scanners and protocols across sites (see Yan et al., 2019 for further details). To manage this multi-scanner variability of data (batch effects), the Correcting Covariance Batch Effects (CovBat) harmonization technique was implemented to correct for site-specific effects directly in the functional connectivity matrices, modeling site as the batch effect and including diagnostic information (MDD or Control) as the variable of interest, as well as sex, age, education years and, head motion (mean framewise displacement) as additional covariates. This method allows the corrected connectivity matrices to retain biologically meaningful patterns while minimizing site-related biases, thus enhancing the comparability of data across multisite studies and improving the robustness of subsequent network-level analyses (A. A. Chen et al., 2022a, 2022b; Wang et al., 2023). The whole harmonization process was conducted using the free CovBat R-package developed by Chen, Beer, et al., (2022).

### Topological Data Analysis: persistent homology

To characterize the topological organization of the functional connectomes, Topological Data Analysis (TDA) was applied using the framework of persistent homology. Persistent homology tracks how topological features of a network, such as connected components, cycles, and other higher-dimensional topological features, emerge, persist, and eventually disappear as a filtration parameter (i.e., sparsity threshold) is gradually varied (Aktas et al., 2019; Centeno et al., 2022). This procedure provides a structured way to describe the topology of a network and enables the detection of higher-order, otherwise hidden patterns of network organization, as well as the identification of their relative stability across scales.

### Filtration and Rips complexes

In topological terms, any weighted network can be described using the Vietoris–Rips complex, denoted as Rips (F, ε), where F represents the nodes and ε is the filtration parameter. Since ε is a distance, the connectivity value needs to be transformed to distance (see below). Here ε is strictly positive and serves as a connectivity threshold: two nodes in F are considered connected if the distance between them is less than ε (Chambers et al., 2010). In a Vietoris-Rips complex (Fig. 2A), a pair of nodes forms a 1-simplex (the analogue of an edge in graph theory) whenever their distance falls below a filtration parameter ε. More generally, a set of k+1 nodes forms a k-simplex whenever all pairwise distances within the set fall below ε (Centeno et al., 2022).

**Fig. 2.**
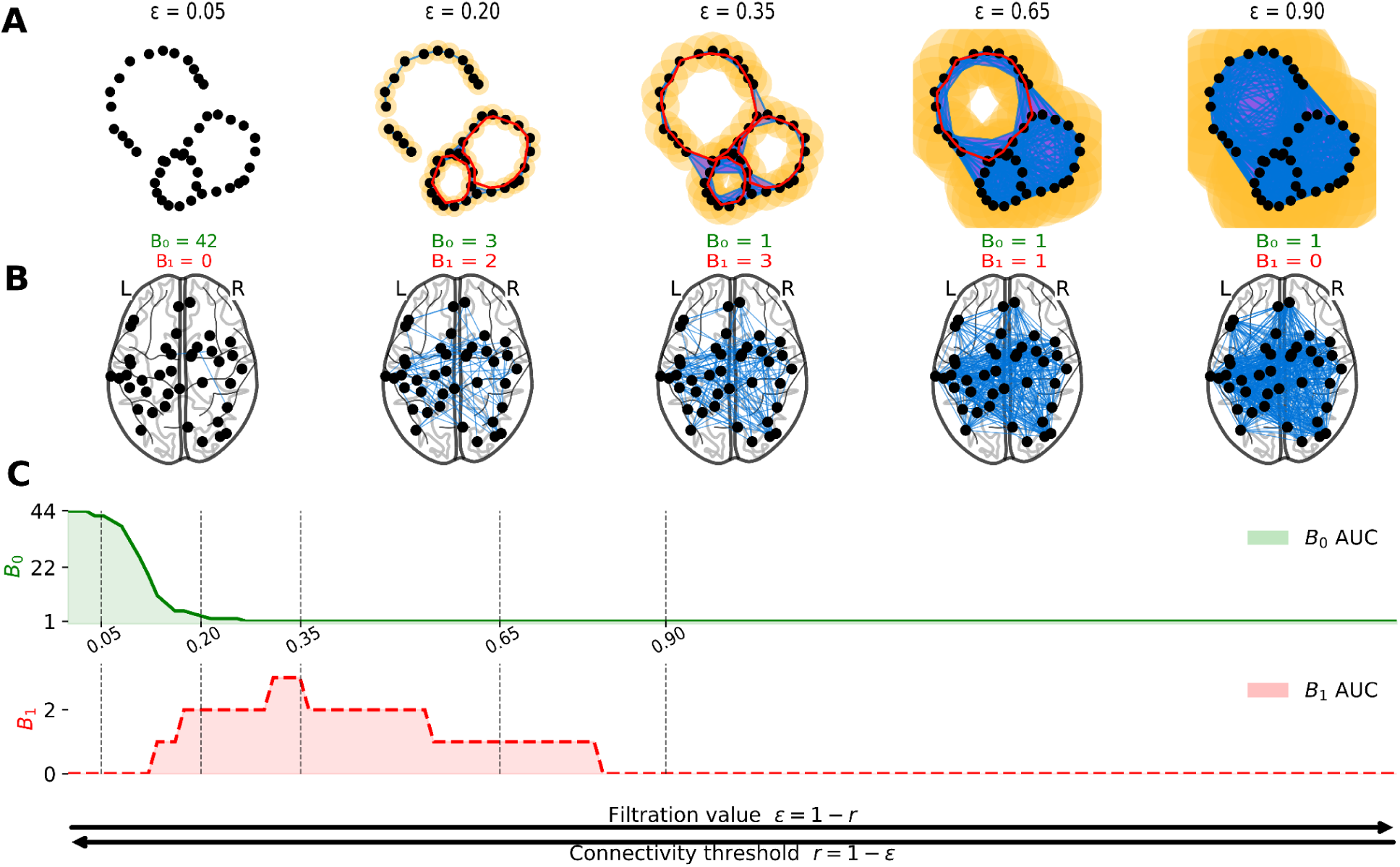
Overview of the persistent homology pipeline applied to functional connectomes. (**A**) Toy example illustrating the construction of Vietoris–Rips complexes across increasing filtration values (ε = 1 - *r*). Nodes are connected whenever their pairwise distance is less than or equal to ε, forming edges (1-simplices; here in blue) and filled triangles (2-simplices; here in purple). For visualization, each node (black dot) is surrounded by balls of radius ε/2 (yellow), such that connections form when these balls intersect. As ε grows, disconnected components merge, leading to changes in the number of connected components (B₀), while non-trivial cycles emerge or vanish, leading to changes in B₁. For visualization purposes, non-trivial cycles surrounding one-dimensional holes are highlighted in red (when the corresponding cycle persists in that simplicial complex). (**B**) Illustration of the corresponding anatomical connectivity graphs projected onto the Power-264 coordinates. At each ε, edges are added whenever pairwise distances fall below the filtration threshold, and the resulting topology of the functional connectome is shown. Betti numbers (B₀ and B₁) computed at each ε summarize the number of connected components and one-dimensional holes (cycles) present in the network at that scale. (**C**) Betti curves. The green curve shows B₀ and the red curve shows B₁, with dashed vertical lines marking the filtration values used in Panels **A–B**. The x-axis is expressed both as a distance threshold (ε) and as the equivalent correlation threshold (r = 1 − ε). The shaded regions represent the area under each curve (AUC). All toy examples were calculated in python using the GUDHI package and common plotting packages.

When ε = 0, all nodes (0-simplices) remain isolated as disconnected components. For small ε, only strongly associated pairs of nodes form 1-simplices, yielding several disconnected components. As ε increases, the complex becomes progressively denser: additional edges appear, then filled triangles (2-simplices), and eventually higher-dimensional simplices (k-simplices). This defines a nested sequence of Vietoris-Rips complexes that constitutes a filtration, which forms the basis of persistent homology (Edelsbrunner and Harer, 2010).

Topological features such as connected components and cycles may appear or disappear throughout the filtration. Persistent homology characterizes this process by tracking the birth and death of such features as a function of ε, as the filtration parameter varies, identifying the values or scales over which each feature exists and providing a concise, multi-scale description of the evolving topology. Each topological feature can be represented by a homology generator that reflects its essential structure across the filtration; for example, in dimension one, generators correspond to non-trivial cycles that capture evolving 1-dimensional topological holes. The birth and death of topological features give rise to the notion of persistence, defined as the difference between the ε value at which a feature appears (birth) and the ε value at which it disappears (death), thereby quantifying its relevance across scales (Centeno et al., 2022).

In our framework, distances were defined as d(xᵢ, x◻) = 1 - r(xᵢ, x◻), where r denotes the Pearson correlation between nodes xᵢ and x◻. Thus, ε represents a distance threshold, and increasing ε corresponds to lowering the traditional correlation threshold used in network neuroscience. This dual interpretation is illustrated in Fig. 2C. In practice, a Vietoris-Rips filtration was constructed for each subject by varying ε across its full theoretical range (0 ≤ ε ≤ 2), as implied by the distance transformation described above. A representative example of this filtration process applied to a real subject is shown in Supplementary Fig. S3.

### Betti numbers

The zeroth Betti number (B₀) counts the number of connected components (0-dimensional holes) in the Vietoris-Rips complex, meaning either isolated nodes or groups of nodes that are internally connected but disconnected from the rest of the complex (Fig. 2A). In contrast, the first Betti number (B₁) counts the number of non-trivial cycles, meaning closed paths (highlighted in red in Fig. 2A) that have not yet been filled in by 2-simplices (triangles). These cycles define the boundaries of 1-dimensional topological holes and therefore provide a topological description of higher-order network organization (Aktas et al., 2019; Centeno et al., 2022).

### Betti curves and area-under-the-curve (AUC)

The evolution of the brain topology was summarized using Betti curves; these were computed for each subject using the TDA R-package (Fasy et al., 2014). Betti curves reflect how many components or non-trivial cycles exist at each ε value (Fig. 2C). As ε increases, B₀ decreases monotonically: disconnected nodes merge into components, and the curve moves from its maximal value at ε = 0 toward a constant value of one once the entire complex becomes a single connected component. In contrast, the B₁ curve follows a characteristic rise-and-fall pattern: it begins at zero, increases as new cycles appear across the filtration, and ultimately returns to zero once those cycles are filled in by higher-dimensional simplices (Edelsbrunner and Harer, 2010; Otter et al., 2017).

To obtain a single global descriptor of each subject’s topology, the area under each Betti curve (AUC; see shaded regions in Fig. 2C) was calculated. The AUC has emerged as a stable and informative metric capable of capturing group-level differences in functional connectivity (Das et al., 2023; Gracia-Tabuenca et al., 2023, 2020). The area under the B₀ and B₁ curves (B₀ AUC and B₁ AUC) were used as the primary outcomes to assess whether global topological organization differs between groups and whether such differences relate to clinically relevant heterogeneity in MDD.

### Interpretation of B₀ AUC

In network settings, the edges responsible for decreasing the B₀ value throughout the filtration are mathematically equivalent to those forming a spanning tree of the network (Chung et al., 2019, 2015; Lee et al., 2012). Depending on whether the filtration is expressed in terms of connection strengths or distances, this structure corresponds to a maximum spanning tree (Chung et al., 2024) or, equivalently, a minimum spanning tree under the distance transformation d = 1 − r (Ryu et al., 2023). Because this spanning-tree structure connects all nodes through the strongest available relationships while avoiding redundant cycles, it has been interpreted as the backbone supporting large-scale integration and information flow across the connectome (Ryu et al., 2023). Accordingly, B₀ AUC quantifies the overall integration profile across filtration values, capturing how disconnected components progressively merge into a single connected structure. Because B₀ is determined by backbone-forming edges, B₀ AUC provides a multiscale summary of how the backbone structure emerges as initially fragmented regions become integrated into a globally connected network, and may therefore be interpreted as an index of multiscale backbone integration.

### Interpretation of B₁ AUC

While B₀ reflects the emergence of network integration, B₁ captures higher-order topological structure that cannot be reduced to pairwise connectivity contributions alone. The cycles identified by B₁ define the boundaries of one-dimensional topological holes (see Figure 2A), allowing the characterization of organizational features that emerge from the collective arrangement of multiple connections rather than from changes in individual edges (Centeno et al., 2022). In turn, variation in B₁ AUC may arise from differences in either the prevalence of hole-defining cycles, the persistence of those cycles across scales, or both. Accordingly, B₁ AUC can be interpreted as an index of hole-defining cycle burden, reflecting the overall prevalence and persistence of these topological configurations.

### Decomposition of B₁ AUC

Given B₁ AUC depends jointly on the number of cycles and their persistence, it was further decomposed into two components: (i) the total number of B₁ cycles per subject, and (ii) their mean persistence (mean lifespan across ε). This decomposition was used to disentangle the relative contributions of cycle abundance and persistence to group-level differences in B₁-related topology, allowing us to determine whether between-group differences in B₁ AUC were driven primarily by a greater number of cycles or by cycles persisting longer across the filtration. From and interpretative perspective, B₁ cycles can be understood as boundaries of topological holes, capturing patterns of organization in which the boundary forms a closed cycle of connections while the interior remains relatively sparsely connected and is not filled by higher-dimensional simplices. Accordingly, a higher number of cycles may reflect a greater prevalence of such higher-order configurations within the network, whereas persistence informs the extent to which these configurations remain expressed across scales.

### Network anatomical context of B₁-cycles

To explore the specific patterns of large-scale brain organization underlying topological differences, the nodes contributing to each cycle (i.e., B₁ homology generator) were localized using Power-264 atlas coordinates, categorizing cycles as intra- vs. inter-network, according to whether they involved nodes within the same functional network or across different networks.

### Network-set motif analysis of inter-network B₁ cycles

To further characterize the organization of inter-network cycles, each B₁ cycle ( i.e., homology generator) was annotated according to the canonical Power-264 networks to which its constituent nodes belonged. This allowed us to identify the particular network configurations underlying inter-network cycles, rather than only determining whether they spanned more than one network. Accordingly, for every cycle a network-set motif was defined as the unique set of resting-state networks spanned by its nodes (e.g., FPN–DMN, DMN–SAL–VIS), such that motifs represent the combination of networks involved, regardless of how many nodes belonged to each network. For this network-set motif analysis, only motifs involving more than one network (set size > 1) were considered, as the aim of this analysis was to specifically characterize cycles spanning multiple networks.

To examine how specific inter-network configurations contribute to group differences in B₁ cycles, we defined a binary presence indicator for each motif in each subject (presence = at least one cycle expressed that motif; absence = otherwise). Group-level motif prevalence was then computed as the proportion of participants in whom the motif was present within each diagnostic group. A motif-wise difference in prevalence was defined as:

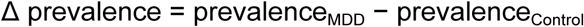

which served as a descriptive index summarizing relative enrichment of each motif between groups. All motifs were retained for descriptive visualization of the distribution of Δ prevalence values. To obtain a global profile of single-network involvement, all motifs with Δ prevalence > 0 were decomposed into their constituent networks and the frequency of each network was computed as the percentage of motifs in which it appeared.

To assess the consistency of these descriptive patterns to the influence of low-persistence cycles, the entire motif-construction and prevalence-estimation procedure were repeated after restricting each subject’s cycles to their 50% most persistent cycles. The resulting single-network involvement profiles were qualitatively similar; these sensitivity analyses are reported in the Supplementary Materials (Supplementary Fig. S3). Finally, for visualization purposes only, the extreme positive tail of the Δ prevalence distribution was identified by selecting the top 1% of motif-wise Δ prevalence values. This step provides an illustrative example of the motifs most strongly biased toward MDD and does not constitute an inferential procedure.

### Statistical Analysis

After deriving topological descriptors from persistent homology, four primary outcomes were analyzed: B₀ AUC, B₁ AUC, B₁ cycle counts, and mean B₁ cycle persistence. Their associations with diagnosis (Equation 1), illness course (Equation 2), and current medication status (Equation 3) were examined in the primary analyses. Depending on the distributional properties of each outcome, different regression frameworks were used: multiple linear regressions (MLRs) for B₀ AUC, Gamma generalized linear models (GLMs) with a log link for B₁ AUC and mean persistence, and negative binomial GLMs for B₁ cycle counts. In all models, the clinical factor of interest served as the primary predictor (see Table 1 of sample composition for group definitions), while age, sex (binary coded), education years, and head motion (mean framewise displacement, FD) were included as covariates.

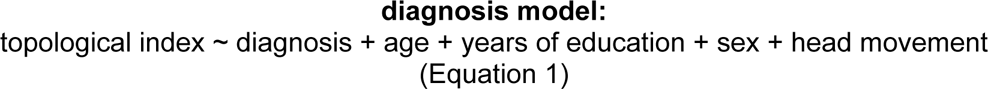

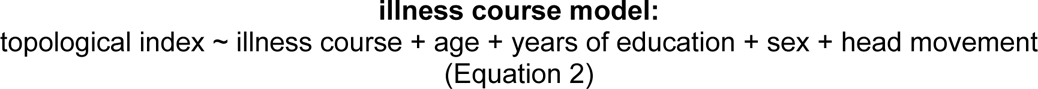

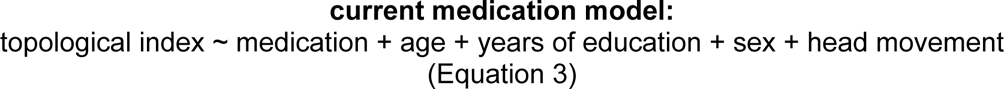

Secondary analyses were restricted to participants with MDD within the recurrence-defined subsample and examined whether topological indices B₀ AUC and B₁ AUC were associated with illness duration or symptom severity (HAMD score). These models followed the same covariate structure as the primary analysis, with illness duration or HAMD score entered as the predictor of interest.

To ensure robust inference, in all models, a bootstrap resampling procedure was used to compute *p*-values for group comparisons, thereby reducing the impact of potential violations of model assumptions. Specifically, 10,000 bootstrap samples were generated by resampling the original dataset with replacement, maintaining the same sample size. For each bootstrap sample, the model was refitted and the group differences of interest were re-estimated accordingly (e.g., between diagnostic groups or recurrence levels); *p*-values were obtained as the proportion of bootstrap estimates exceeding the observed difference in absolute value. To account for multiple comparisons, the Benjamini–Hochberg false discovery rate (FDR) correction was applied. In practice, all post-hoc contrasts were computed on the link scale using two-sided bootstrap tests (10,000 repetitions; FDR-adjusted).

Effect sizes were estimated according to the model type. For MLRs, Cohen’s d was computed as the ratio between the estimated regression coefficient for the condition of interest and the residual standard deviation 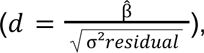 where β̂ is the estimated regression coefficient for the variable of interest and σ²_residual_ is the residual variance. This approach, recently recommended for regression models with covariates (Groß and Möller, 2024, 2023), facilitates interpretation of the condition’s effect relative to unexplained variability. For GLMs with Gamma error distribution and log link, the estimated β coefficients were exponentiated to obtain fold-change ratios on the response scale (Hardin and Hilbe, 2018). This transformation is appropriate because the log link models the logarithm of the outcome as a linear combination of predictors, such that differences on the log scale correspond to multiplicative changes in the original outcome scale. Finally, for negative binomial GLMs the coefficients were likewise exponentiated and reported as incidence rate ratios (IRRs) (Hardin and Hilbe, 2018). All statistical analyses were performed in R.

To facilitate interpretation of the overall analytic workflow, Fig. 3 provides a schematic overview of the complete processing pipeline, summarizing the transformation from resting-state fMRI data to topological descriptors and their subsequent statistical analysis.

**Fig. 3.**
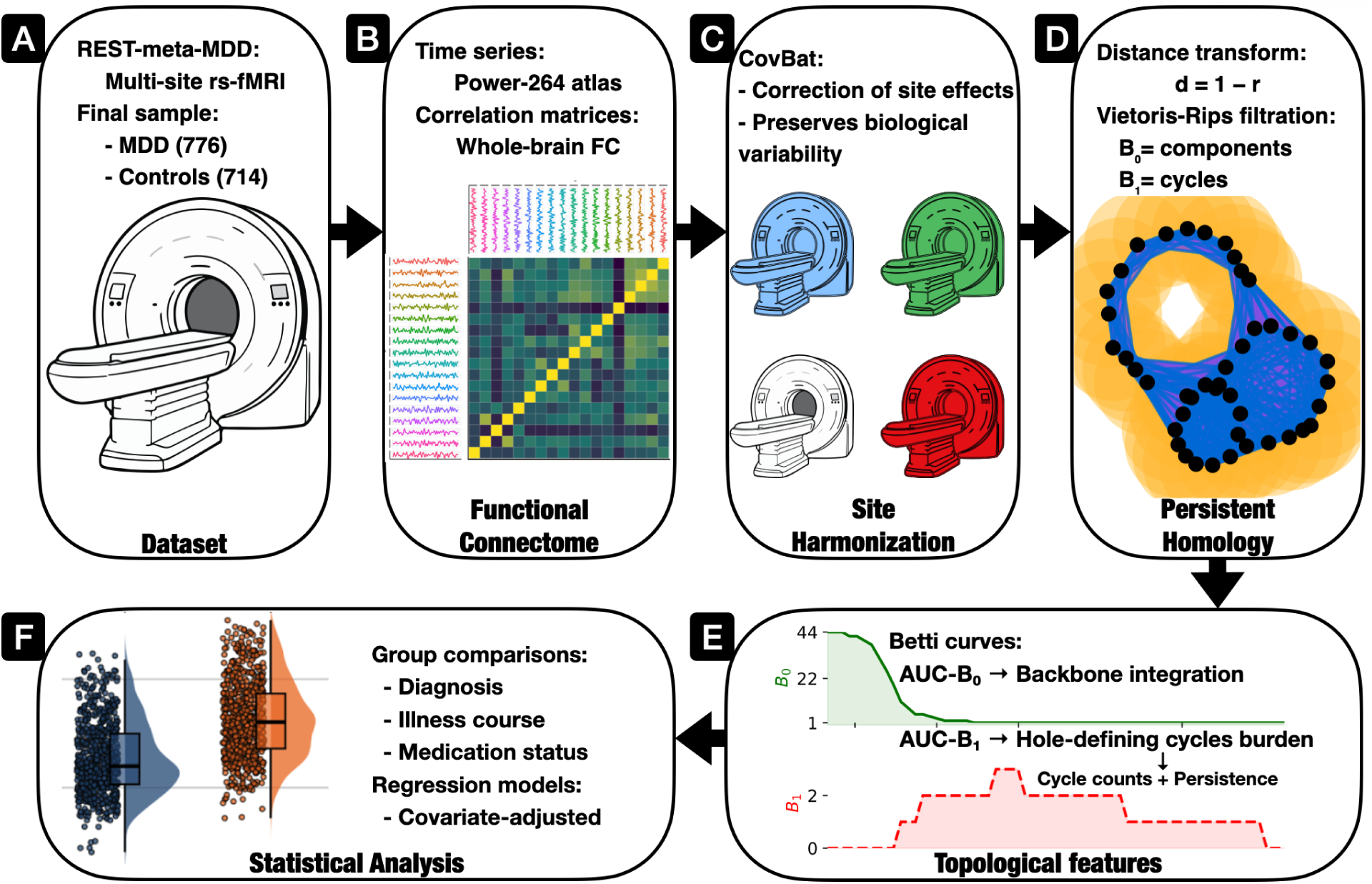
Overview of the analytical pipeline. Panels **A–D** depict preprocessing and topological feature extraction, panel **E** summarizes the resulting topological descriptors, and panel **F** shows the primary statistical analyses. **(A)** Resting-state fMRI data were obtained from the REST-meta-MDD consortium after quality-control and sample selection. **(B)** Construction of subject-level functional connectomes from Power-264 regional time series. **(C)** Site harmonization using CovBat. **(D)** Transformation of harmonized correlation matrices into distance matrices (d = 1 − r) and construction of Vietoris–Rips filtrations to characterize connected components (B₀) and cycles (B₁). **(E)** Extraction of Betti curves and their areas under the curve (AUC), yielding indices of multiscale backbone integration (B₀ AUC) and hole-defining cycle burden (B₁ AUC), with additional decomposition of B₁ into cycle counts and persistence. **(F)** Regression statistical models relating topological metrics to diagnosis, illness course, medication status, and other clinical variables, and adjusted by relevant covariates.

### Reproducibility and code availability

All custom scripts used in this study are publicly available at: https://github.com/soundingreen/TDA-MDD_DIRECT.git. The repository includes the complete analytical workflow, including participant selection and quality-control procedures, construction of analytic subsamples, functional-connectome generation, CovBat harmonization, persistent homology computations, Betti-curve and AUC extraction, B₁-cycle decomposition analyses, network-set motif analyses, statistical modeling, and figure generation. Analyses were performed in R and some figures were created using Python. Access to the original REST-meta-MDD dataset is governed by the DIRECT consortium.

## Results

### Whole-brain topology

In the full sample with complete data for the P264 atlas (**n = 1,490**), consistent group differences were found in the topological indices B₀ AUC and B₁ AUC, reflecting distinct patterns of whole-brain network organization in MDD. For B₀ AUC (Fig. 4, Panel A), individuals with MDD showed significantly higher values than controls (β = 1.66, Cohen’s d = 0.19, FDR-*p* < 0.001, adjusted R² = 0.16), suggesting a more gradual integration process at the whole-brain level in MDD. A similar pattern emerged for B₁ AUC (Fig. 4, Panel B), where MDD participants displayed higher B₁ AUC values than controls (β = 0.05, exp(β) = 1.05, FDR-*p* <0.001, Nagelkerke’s R² = 0.05), consistent with a greater prevalence of hole-defining cycles, greater persistence of those cycles across scales, or both. In a subset of participants with available data on episode recurrence (**n = 1,296**), further analysis (Fig. 4C) revealed that both single-episode (MDD-SE) and recurrent (MDD-R) depression groups had significantly elevated B₀ AUC compared to controls (MDD-SE: β = 1.47, Cohen’s d = 0.16, FDR-*p* < 0.05; MDD-R: β = 1.95, Cohen’s d = 0.22, FDR-*p* < 0.05), with no significant difference between these two subgroups (β = 0.48, FDR-*p* = 0.59). In contrast, B₁ AUC (Fig. 4D) exhibited a clear gradient across illness course: MDD-SE (β = 0.04, FDR-*p* < 0.01, exp(β) = 1.04) and MDD-R showed higher values (β = 0.011, exp(β) = 1.11, FDR-*p* < 0.001, Nagelkerke’s R²= 0.055) compared to controls, with MDD-R also showing significantly higher B₁ AUC than MDD-SE (β =0.07, exp(β) = 1.08, FDR-*p* < 0.001), indicating that higher-order topological differences become more pronounced with illness progression.

**Fig. 4.**
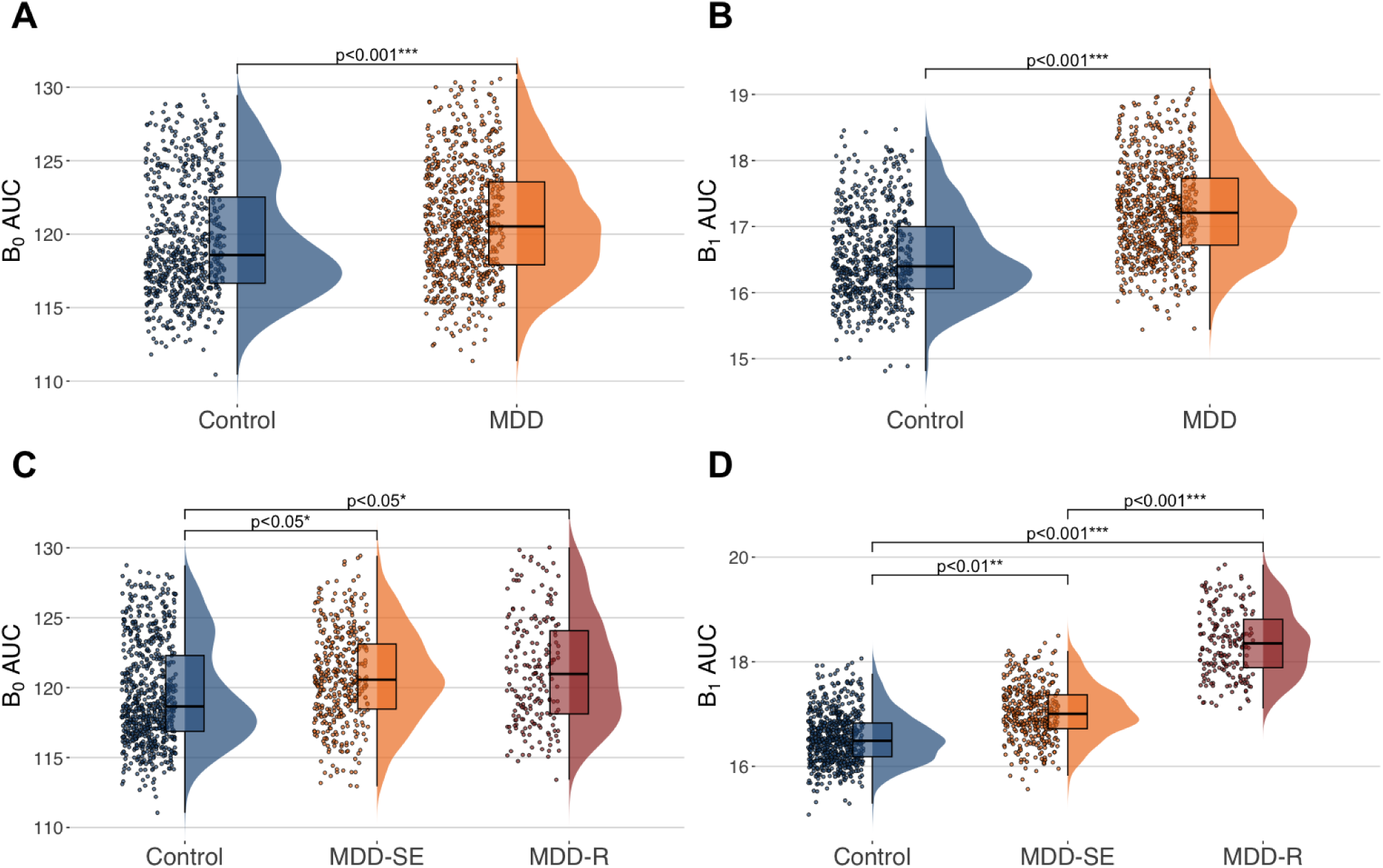
Whole-brain topological differences between controls and MDD. **(A)** Individuals with MDD (orange) showed significantly higher whole-brain Betti-0 AUC values than controls (blue), indicating altered integration of functional components. **(B)** A similar pattern was observed for Betti-1 AUC, with MDD participants exhibiting an increased index. **(C)** Both single-episode (MDD-SE, orange) and recurrent (MDD-R, burgundy) participants showed elevated Betti-0 AUC compared to controls (blue), with no significant difference between subgroups. **(D)** Betti-1 AUC showed a progressive increase from controls to MDD-SE and MDD-R, suggesting topological sensitivity to illness course. Displayed values represent model-predicted estimates; all statistical inferences derive from the original data.

Following the observed topological differences between recurrence-defined subgroups of MDD, we explored whether such alterations could be related to other clinically relevant variables. Specifically, we assessed whether whole-brain B₀ AUC and B₁ AUC values were associated with either illness duration or symptom severity (HAMD score) in the MDD group within the recurrence-defined subsample. A total of **n = 538** had data on illness duration, and **n = 511** had HAMD scores. Neither clinical variable showed a significant association with topological indices (all *p*-values > 0.05), suggesting that neither illness duration nor current symptom severity accounted for the observed topological changes associated with presence and recurrence of MDD.

Next, it was examined whether current medication status, stratified by illness course, could better account for the observed topological differences in MDD. In the subsample of participants with complete information on both recurrence and medication status (**n = 1,166**), the effect of medication at the time of scanning, stratified by recurrence was assessed as the next step. While B₀ AUC remained unaffected across all medication-defined subgroups (all *p*-values > 0.05), B₁ AUC exhibited significant differences (Fig. 5). Compared to controls, the unmedicated single-episode MDD group (uMDD-SE) showed a significant 7% increase (β = 0.0688, FDR-*p* < 0.01, exp(β) = 1.07, Nagelkerke’s R² = 0.05). The medicated recurrent group (mMDD-R) also exhibited a robust 9% increase (β = 0.0888, exp(β) = 1.09, FDR-*p* < 0.01). In contrast, other subgroups (mMDD-SE and uMDD-R) did not differ significantly from controls after FDR correction. Notably, mMDD-SE showed significantly lower B₁ AUC values than uMDD-SE (β = –0.0863, FDR-*p* < 0.01, exp(β) = 0.92), and mMDD-R differed from mMDD-SE (β = 0.1064, FDR-*p* < 0.01, exp(β) = 1.11), indicating that both illness course and medication jointly influence the topology of the functional connectome.

**Fig. 5.**
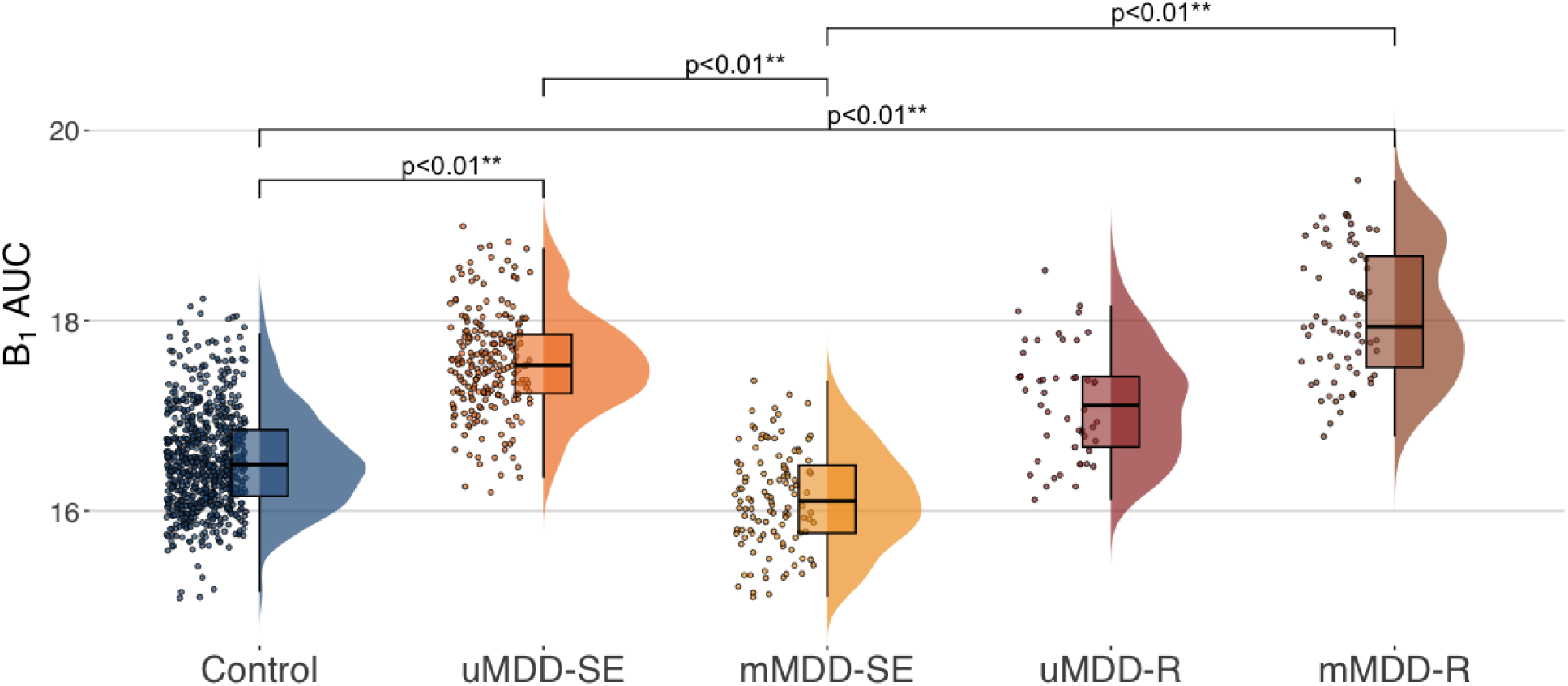
Effects of medication and illness course on whole-brain B₁ AUC. Compared to controls (blue), Betti-1 AUC was significantly higher in the unmedicated single-episode (uMDD-SE, orange) and the medicated recurrent (mMDD-R, brown) groups. No significant differences were found for the medicated single-episode (mMDD-SE, yellow) and unmedicated recurrent (uMDD-R, burgundy) groups. However, Betti-1 AUC was lower in mMDD-SE compared to uMDD-SE, and mMDD-R differed significantly from mMDD-SE, indicating an interaction between recurrence and medication status. Plots depict model-derived predictions; *p*-values and effect sizes are based on fits to the original data.

To pinpoint the source of the B₁ AUC increase, this increase was decomposed into the number of B₁ cycles counts (Fig. 6A) and mean persistence. MDD showed a significant increase in the number of B₁ cycles relative to controls (β = 0.047, FDR-*p* < 0.001; IRR = 1.048, 95% CI [1.028 – 1.069], Nagelkerke’s R² = 0.15), whereas mean persistence did not differ between groups (p > 0.05). Thus, the whole-brain B₁ AUC effect appears to be driven by a greater number of hole-defining cycles rather than by increased cycle persistence. Extending this analysis to illness course (Fig. 6B), both single-episode (MDD-SE; β = 0.039, FDR-*p* < 0.01; IRR = 1.039, 95% CI [1.015–1.064]) and recurrent depression (MDD-R; β = 0.101, FDR-*p* < 0.001; IRR = 1.106,, 95% CI [1.073 – 1.139]) were associated with elevated cycle counts relative to controls, with significantly greater values in MDD-R compared to MDD-SE (β = 0.061, FDR-*p* < 0.001; IRR = 1.064, 95% CI [1.031 – 1.098]). Mean persistence remained comparable across groups (all p> 0.05). Together, these results indicate that differences in B₁ AUC between recurrence-defined subgroups are primarily driven by an increased number of one-dimensional cycles (B₁ cycles) in MDD, particularly in recurrent cases, rather than by differences in cycle persistence.

**Fig. 6.**
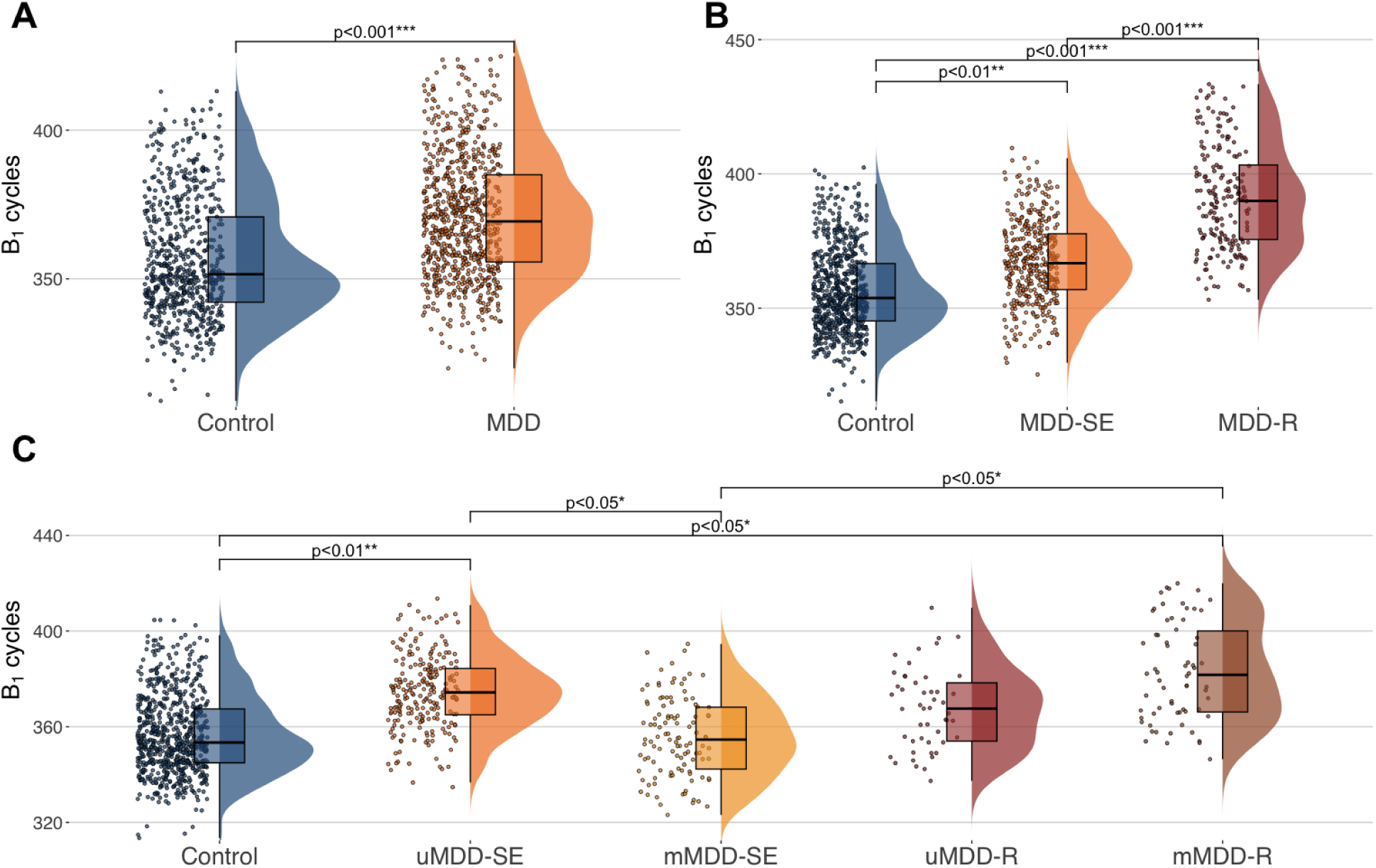
Number of B₁ cycles. **(A)** Individuals with MDD (orange) showed significantly higher B₁ cycle counts than controls (blue), indicating a greater prevalence of hole-defining cycles. **(B)** B₁ cycles showed a progressive increase from controls to MDD-SE and MDD-R, suggesting topological sensitivity to illness course. **(C)** Compared to controls (blue), B₁ cycles was significantly higher in the unmedicated single-episode (uMDD-SE, orange) and the medicated recurrent (mMDD-R, brown) groups. No significant differences were found for the medicated single-episode (mMDD-SE, yellow) and unmedicated recurrent (uMDD-R, burgundy) groups. However, B₁ cycles was lower in mMDD-SE compared to uMDD-SE, and mMDD-R differed significantly from mMDD-SE, indicating an interaction between recurrence and medication status. Displayed values represent model-predicted estimates; all statistical inferences derive from the original data.

Finally when stratifying by medication status (Fig. 6C), unmedicated single-episode MDD group (uMDD-SE) showed significantly higher number of B₁ cycles compared to controls (β = 0.056, FDR-p < 0.01; IRR = 1.058, 95% CI [1.027–1.090], Nagelkerke’s R² = 0.132). The medicated recurrent group (mMDD-R) also exhibited a higher number of B₁ cycles compared to controls (β = 0.066, FDR-p < 0.05; IRR = 1.067, 95% CI [1.023–1.114]). In contrast, other subgroups (mMDD-SE and uMDD-R) did not differ significantly from controls after FDR correction. Notably, mMDD-SE showed significantly lower B₁ AUC values than uMDD-SE (β = -0.059, FDR-p < 0.05; IRR = 0.943, 95% CI [0.906–0.980]), and mMDD-R differed from mMDD-SE (β = 0.067, FDR-p < 0.05; IRR = 1.070, 95% CI [1.018–1.125]), indicating that both illness course and medication jointly influence the topology of the functional connectome. Medication status did not yield significant effects regarding the mean persistence of Betti-1 cycles (all FDR-p > 0.05). These results suggest that the excess of one-dimensional cycles in MDD is most pronounced during untreated first episodes, whereas in recurrent episodes it becomes most evident under pharmacological treatment.

### Network anatomical context of B₁ cycles

To further improve the interpretability of the higher-order topological alterations captured by B₁ cycles (B₁ homology generators), their anatomical distribution was examined across large-scale functional networks. Initially, a distinction was made between intra- and inter-network cycles without considering the specific network identities involved. No significant group differences emerged in the number of intra-network cycles between MDD participants and controls (all FDR-*p* > 0.05). By contrast, inter-network cycles were significantly increased in the MDD group relative to controls (β = 0.051, FDR-*p* < 0.001; IRR = 1.053, 95% CI [1.031, 1.076]; Nagelkerke’s R² = 0.17). Further analyses stratified by illness course revealed no differences in the number of intra-network cycles but both single-episode (MDD-SE; β = 0.044, FDR-*p* < 0.01; IRR = 1.045, 95% CI [1.019–1.071]) and recurrent MDD participants (MDD-R; β = 0.107, FDR-*p* < 0.01; IRR = 1.113, 95% CI [1.078–1.149]) exhibited more inter-network cycles than controls, with the strongest effect in the recurrent group. The MDD-R number of intra-network cycles was also significantly higher than MDD-SE (β = 0.063, FDR-*p* < 0.01; IRR = 1.065, 95% CI [1.030–1.101]).

Furthermore, when stratifying by medication status, unmedicated single-episode MDD group (uMDD-SE) showed significantly higher number of inter-network cycles compared to controls (β = 0.064, FDR-*p* < 0.01; IRR = 1.066, 95% CI [1.032–1.100], Nagelkerke’s R² = 0.149).The medicated recurrent group (mMDD-R) also exhibited a robust increase (β = 0.070, FDR-*p* < 0.05; IRR = 1.072, 95% CI [1.025–1.122]). In contrast, mMDD-SE and uMDD-R did not differ significantly from controls after FDR correction. Notably, mMDD-SE showed significantly lower number of inter-network cycles than uMDD-SE (β = -0.061, FDR-*p* < 0.05; IRR = 0.940, 95% CI [0.900–0.980]), and mMDD-R have higher inter-network cycles than mMDD-SE (β = 0.067, FDR-*p* < 0.05; IRR = 1.070, 95% CI [1.014–1.130]). No differences in the number of intra-network cycles were evident. Altogether, these results suggest that the topological alterations observed in MDD are primarily driven by an increased number of inter-network B₁ cycles across large-scale brain systems rather than within-network organization.

### Motif-based characterization of inter-network B₁ cycles

Finally, to provide an exploratory overview of the functional systems represented in inter-network Betti-1 cycles, each cycle was classified by the set of canonical Power-264 networks spanned by its constituent nodes. Each cycle was therefore assigned to a network-set motif, defined as the unique combination of networks involved (e.g., a cycle including nodes from the default mode (DMN), salience (SAL), and visual (VIS) networks would be labeled as DMN–SAL-VIS, regardless of how many nodes belonged to each network), and motif prevalence was computed as the proportion of participants in each diagnostic group expressing at least one cycle with that configuration.

The distribution of group differences in motif prevalence (Δ prevalence = prevalence_MDD_ − prevalence_Control_) was centered near zero, with a positive tail reflecting motifs more frequently observed in MDD (Fig. 7A). All motifs with Δ prevalence > 0, were decomposed into their constituent networks to obtain a global profile of the networks being involved. This analysis indicated that somatomotor-hand (SMH), default mode (DMN), cingulo-opercular (CON), uncertain/heteromodal (UNC), and visual (VIS) networks were the most frequently represented among inter-network motifs enriched in MDD (Fig. 7B).

**Fig. 7.**
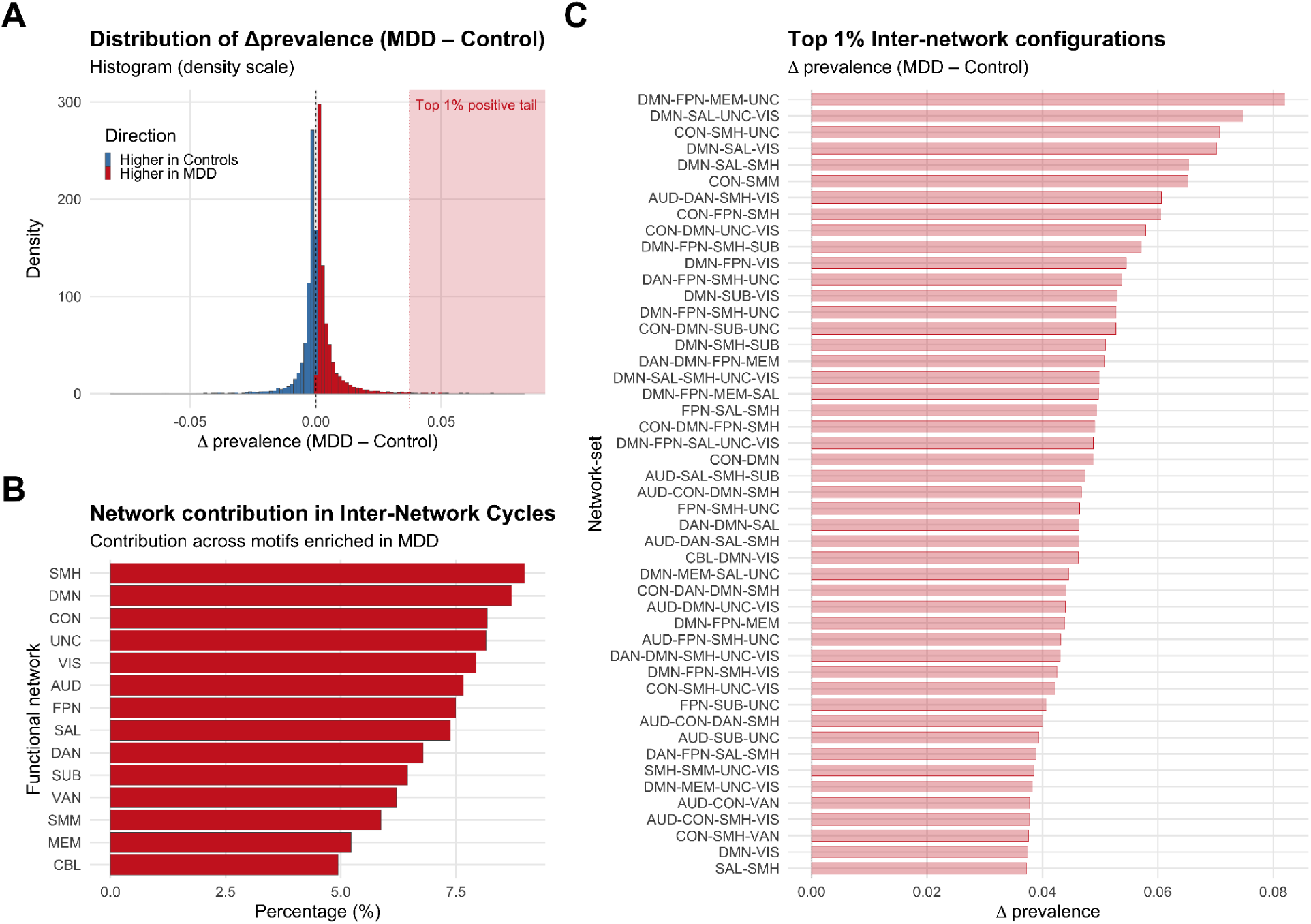
Network-set motifs of inter-network B₁ cycles. **(A)** Distribution of between-group differences in motif prevalence (Δprevalence = MDD – Control) across all inter-network motifs. The extreme positive tail (top 1%) is highlighted in red. **(B)** Decomposition of all motifs with higher prevalence in MDD (Δprevalence > 0) into their constituent single-networks, showing the total contribution of each functional system. Bars indicate the percentage of these motifs that include each canonical Power-264 network. The most represented networks were somatomotor-hand (SMH), default mode (DMN), cingulo-opercular (CON), uncertain/heteromodal (UNC), and visual (VIS). **(C)** Motifs in the extreme positive tail (top 1% of Δprevalence values), shown as illustrative examples of strongly MDD-biased inter-network configurations. This panel is descriptive and not inferential; motif identities may vary with persistence thresholds, whereas the single-network profile in panel B remains consistent. Network abbreviations (Power-264 atlas): SMH = Somatomotor-Hand; SMM = Somatomotor-Mouth; CON = Cingulo-opercular; DMN = Default Mode; FPN = Fronto-parietal; SAL = Salience; DAN = Dorsal Attention; VAN = Ventral Attention; VIS = Visual; AUD = Auditory; MEM = Memory Retrieval; SUB = Subcortical; CBL = Cerebellar; UNC = Uncertain.

As an illustrative example, the small set of motifs falling within the extreme positive tail (top 1% of Δ prevalence values) was highlighted, corresponding to the most strongly MDD-skewed configurations (Fig. 7C). While the precise motif identities in this tail vary with persistence thresholds, reflecting the combinatorial nature of network-set labels, the overall single-network profile remained stable. A sensitivity analysis restricted to the 50% most persistent cycles per subject yielded a qualitatively similar ordering of network involvement (Supplementary Fig. S4), indicating that the prominence of SMH, DMN, CON, UNC, and VIS in MDD-enriched motifs is robust to reasonable variations in filtering. Taken together, these findings indicate that inter-network B₁ cycles in MDD more frequently involve specific large-scale systems, particularly SMH, DMN, CON, UNC, and VIS, highlighting a structured pattern of network involvement.

## Discussion

In this large multisite study, we provide a multiscale characterization of the whole-brain functional connectome in MDD using topological descriptors derived from persistent homology, while also assessing how these patterns vary along clinically relevant dimensions of heterogeneity, namely illness course and current medication status.

Across the full cohort, individuals with MDD showed systematic differences in whole-brain topology compared with controls, reflected in both zeroth- and first-order Betti summaries. These findings support the notion that depression involves large-scale disturbances in functional brain connectivity rather than focal abnormalities and extend prior rs-fMRI work by characterizing connectivity across a multiscale topological representation. Importantly, however, these alterations were not uniformly reflected across topological descriptors. Instead, different descriptors appeared sensitive to different aspects of the disorder, suggesting that different facets of whole-brain topology may capture distinct dimensions of MDD rather than a single topological signature. While both B₀ and B₁ distinguished MDD from controls, only B₁ varied systematically according to illness course and medication status. This dissociation suggests that different topological descriptors capture distinct levels of organization and may reflect different aspects of disease-related heterogeneity.

From a topological perspective, B₀ AUC captures the global integration profile of the network across sparsity scales (Das et al., 2023; Lee et al., 2012; Ryu et al., 2023). Because changes in B₀ only occur when a link connects previously disconnected components, B₀ is determined by the subset of connections responsible for forming the connectivity backbone of the network, a structure that has been interpreted as critical for supporting efficient information flow (Chung et al., 2024, 2023; Ryu et al., 2023). Unlike conventional connectivity measures, B₀ AUC does not quantify connectivity strength at a single threshold, nor does it directly reflect conventional notions of hyper- or hypoconnectivity. Rather, it summarizes the efficiency of the progression from network fragmentation to global connectedness across scales. Within this framework, elevated B₀ AUC values indicate a more gradual (less efficient) transition toward global integration, suggesting that the connectome remains fragmented across a wider range of connectivity thresholds. Conversely, lower B₀ AUC values indicate a more rapid transition toward global integration through earlier backbone formation.

B₀ AUC was elevated in MDD but showed no relationship with illness duration, symptom severity, illness course, or medication status. The stability of this effect across clinical dimensions suggests that altered global integration may represent a relatively stable disorder-level characteristic of MDD rather than a marker of episode accumulation or current clinical state. This pattern is consistent with reduced large-scale cohesion and greater functional segregation rather than efficient global coordination (Ryu et al., 2023), and more broadly aligns with prior rs-fMRI literature describing MDD as involving widespread alterations in large-scale brain connectivity (Javaheripour et al., 2021; Kaiser et al., 2015; Li et al., 2018; Mulders et al., 2015).

In contrast, B₁-associated topological descriptors revealed clear sensitivity to clinically relevant heterogeneity. B₁ AUC increased progressively from controls to single-episode and recurrent MDD, indicating that topological alterations captured by B₁ are not uniformly expressed across the disorder, but instead appear to track differences in illness course. However, this sensitivity was not restricted to illness course, as medication status also modulated B₁ AUC. Specifically elevated B₁ AUC values were most prominent in unmedicated single-episode MDD and medicated recurrent MDD, whereas other subgroups did not differ from controls. Notably, B₁-related sensitivity did not extend to current symptom severity, as no association was observed between B₁-derived descriptors and HAMD scores. Thus, the overall pattern captured by B₁ descriptors is characterized by a graded increase across illness course, modulation by medication status, and apparent independence from current symptom burden. Such a profile suggests that the alterations captured by B₁ may be more closely related to longer-term disease trajectory than to momentary symptom severity. The dissociation between symptom severity, illness course, and medication status observed here is compatible with frameworks proposing that first-onset and recurrent depression are only partially overlapping phenomena and may involve distinct biological processes across illness progression (Buckman et al., 2018; Van Loo et al., 2018). Moreover, the joint sensitivity of B₁ descriptors to illness course and medication status may prove relevant for understanding how distinct clinical trajectories are reflected in the brain connectome across the course of depression.

From a topological perspective, B₁ captures the presence of topological holes within the connectome. Decomposition analyses indicated that the observed B₁ increase was driven primarily by a greater number of cycles rather than increased cycle persistence. Consistently, the absence of differences in mean persistence suggests that illness-course effects are not driven by the stabilization of specific topological holes once they emerge, but rather by an increase in the number of hole-defining configurations throughout the connectome. Importantly, the observed increase in B₁ cycles should not be interpreted as evidence of greater redundancy. Instead, it suggests a greater prevalence of configurations in which distributed brain regions are sufficiently connected to form cycle boundaries, yet insufficiently connected to eliminate the corresponding topological holes through additional links (Lee et al., 2014). In this sense, the increase in cycle counts may reflect a greater prevalence of partially integrated large-scale configurations rather than more cross-system redundancy. Previous works similarly associate higher B₁ values with the presence of more hole-defining configurations and less densely connected local organization, whereas reductions in B₁ are associated with a greater completion of local connectivity patterns (Gracia-Tabuenca et al., 2023; Lee et al., 2014). Configurations of this kind are inherently non-pairwise and emerge from the coordinated organization of multiple connections. Persistent homology is particularly well suited to characterize such structures because it tracks their expression across connectivity scales, while preserving information about their higher-order topological organization, thereby complementing conventional pairwise and single-threshold descriptions of functional connectivity (Gracia-Tabuenca et al., 2023; Lee, 2019; Lee et al., 2012).

Anatomical decomposition of B₁ cycles provides further insight into the network-level substrate of these alterations. The increase in cycle counts was driven predominantly by inter-network rather than intra-network cycles, indicating that the heterogeneity captured by B₁ is expressed primarily through interactions among large-scale brain systems rather than through reorganization confined within individual networks. Accordingly, differences associated with illness course and medication status may manifest at the level of cross-network topological holes, suggesting that relevant variation captured by B₁ is concentrated in how distributed functional systems coordinate with one another across the connectome.

On a more exploratory level, the networks most frequently represented in inter-network cycles more prevalent in MDD included the default mode, cingulo-opercular, somatomotor, visual, and heteromodal systems. Rather than indicating a specific or unique “depression motif,” these findings delineate a coarse anatomical context in which topological cycle structures are likely to emerge. This organization is broadly consistent with rs-fMRI work describing depression as involving disrupted coordination among multiple large-scale functional systems, rather than confined to single networks (Javaheripour et al., 2021; Kaiser et al., 2015; Mulders et al., 2015; Zhang et al., 2025). In particular, the prominence of default mode and cingulo-opercular systems aligns with prior evidence implicating disruption in internally oriented processing and cognitive control networks in depression (Duran and Miller, 2020; Menon, 2011; Zhang et al., 2024a; Zheng et al., 2015), while the involvement of sensorimotor and visual systems suggests that such alterations may extend beyond traditionally emphasized affective and control-related circuits (Ray et al., 2021; Zhang et al., 2024b; Zhu et al., 2025).

Viewed together, the progressive increase in B₁ alterations across illness course, their modulation by medication status, and their predominance within inter-network configurations suggest that clinically relevant heterogeneity in MDD is expressed through alterations in large-scale cross-network coordination and is unlikely to be governed by a single network phenotype. The prominence of cross-network alterations across illness course resonates with theoretical frameworks emphasizing distributed vulnerability and stress sensitization across illness course (Monroe and Harkness, 2005), as well as with empirical evidence showing progressive cognitive and functional burden in recurrent depression (Ahern et al., 2025; Semkovska et al., 2019). Nevertheless, any direct relationship between these clinical consequences and the topological alterations observed here remains speculative, as measures of cognitive performance and functional impairment were not available in the present dataset.

However, the medication-related findings should not be interpreted as evidence of treatment effects per se. Medication status in this dataset likely captures complex heterogeneous trajectories of treatment exposure and illness course, which cannot be disentangled with the available information in the dataset. This limitation is particularly relevant given that illness trajectories vary substantially across individuals with MDD, and multiple factors (including symptom profile, treatment response, treatment duration, medication type, episode count, and other clinical variables) may contribute to the observed subgroup differences (Buch and Liston, 2021). In turn, medication-related effects should be interpreted as indexing clinically relevant heterogeneity rather than causal treatment-related changes in the connectome.

Several limitations should be acknowledged. First, the cross-sectional design precludes causal inference regarding illness progression or treatment effects. Longitudinal studies will be required to determine whether topological alterations precede, accompany, or follow clinical changes. Second, medication status was treated as a categorical variable. More detailed information regarding treatment exposure and clinical trajectories was not available in the current dataset limiting our ability to disentangle the sources of medication-related heterogeneity. Future studies incorporating richer treatment metadata will be necessary to clarify these effects. Finally, while the present analyses focused on whole-brain topology, future work could further integrate topological descriptors with graph-theoretical measures and develop more refined localization approaches for topological features, helping bridge global network organization and regionally specific dysfunction.

In summary, the present study demonstrates that MDD is associated with alterations in whole-brain functional organization that are detectable at multiple topological scales. At the level of B₀ depression is characterized by a less efficient multiscale integration process consistent with reduced large-scale cohesion and a more gradual emergence of the connectivity backbone underlying global integration. At the level of B₁ depression is characterized by greater prevalence of cross-network topological holes, reflected in an increased number of inter-network cycles rather than increased cycle persistence. Importantly, these B₁-related topological alterations varied systematically with illness course and medication status, whereas B₀ remained comparatively stable. Collectively, these findings suggest that zeroth- and first-order topological descriptors capture complementary aspects of large-scale brain organization in MDD. While B₀ may reflect relatively stable alterations in global integration, B₁ appears more sensitive to clinically meaningful heterogeneity associated with disease trajectory and treatment context.

By providing a non-pairwise and multiscale characterization of the connectome, persistent homology offers a complementary framework for studying the distributed neurobiology of depression and for understanding how clinically relevant heterogeneity is expressed within large-scale network organization.

## Acknowledgements

Javier F. Castilla-Jiménez is a doctoral student of the Programa de Doctorado en Ciencias Biomédicas at Universidad Nacional Autónoma de México (UNAM) and has received a fellowship (No. 1310997) from Secretaría de Ciencias, Humanidades, Tecnología e Innovación (SECIHTI, formerly CONAHCYT). Juan Carlos Díaz-Patiño was awarded a postdoctoral fellowship from UNAM’s Dirección General de Asuntos del Personal Académico (DGAPA). Sarael Alcauter was awarded a Grant (IN208622) from UNAM’s Programa de Apoyo a Proyectos de Investigación e Innovación Tecnológica (PAPIIT). We thank Nuri Aranda López from PDCB for her administrative support. We thank Leopoldo González Santos for his technical support. We thank Omar González Hernández, Ramon Martinez Olvera, Moisés Mendoza Baltazar, and Maria Eugenia Rosas Alatorre, for their computer systems assistance.

## Supplementary

**Supplementary Fig. S1.**
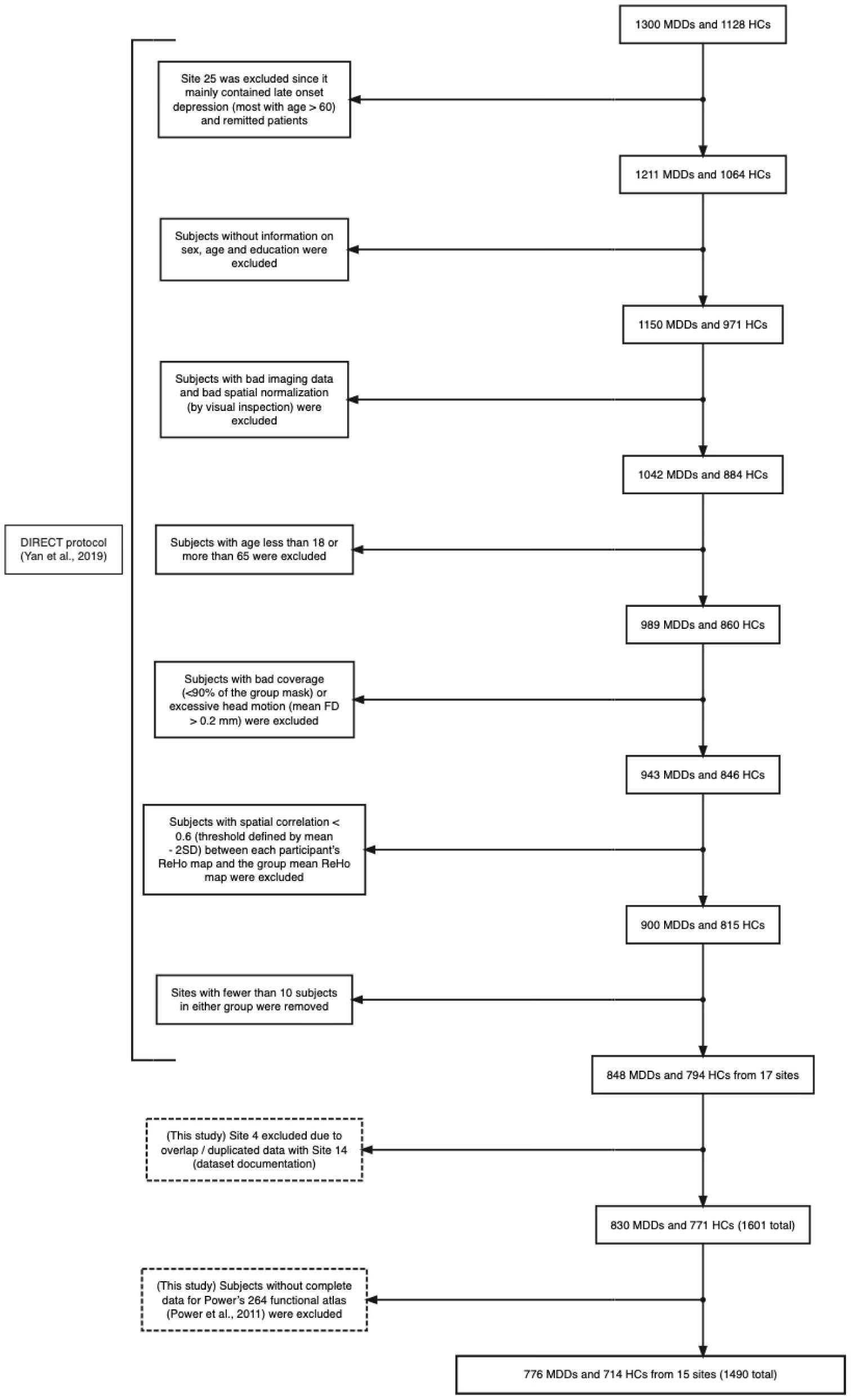
COHORT flow chart for sample selection.

**Supplementary Fig. S2.**
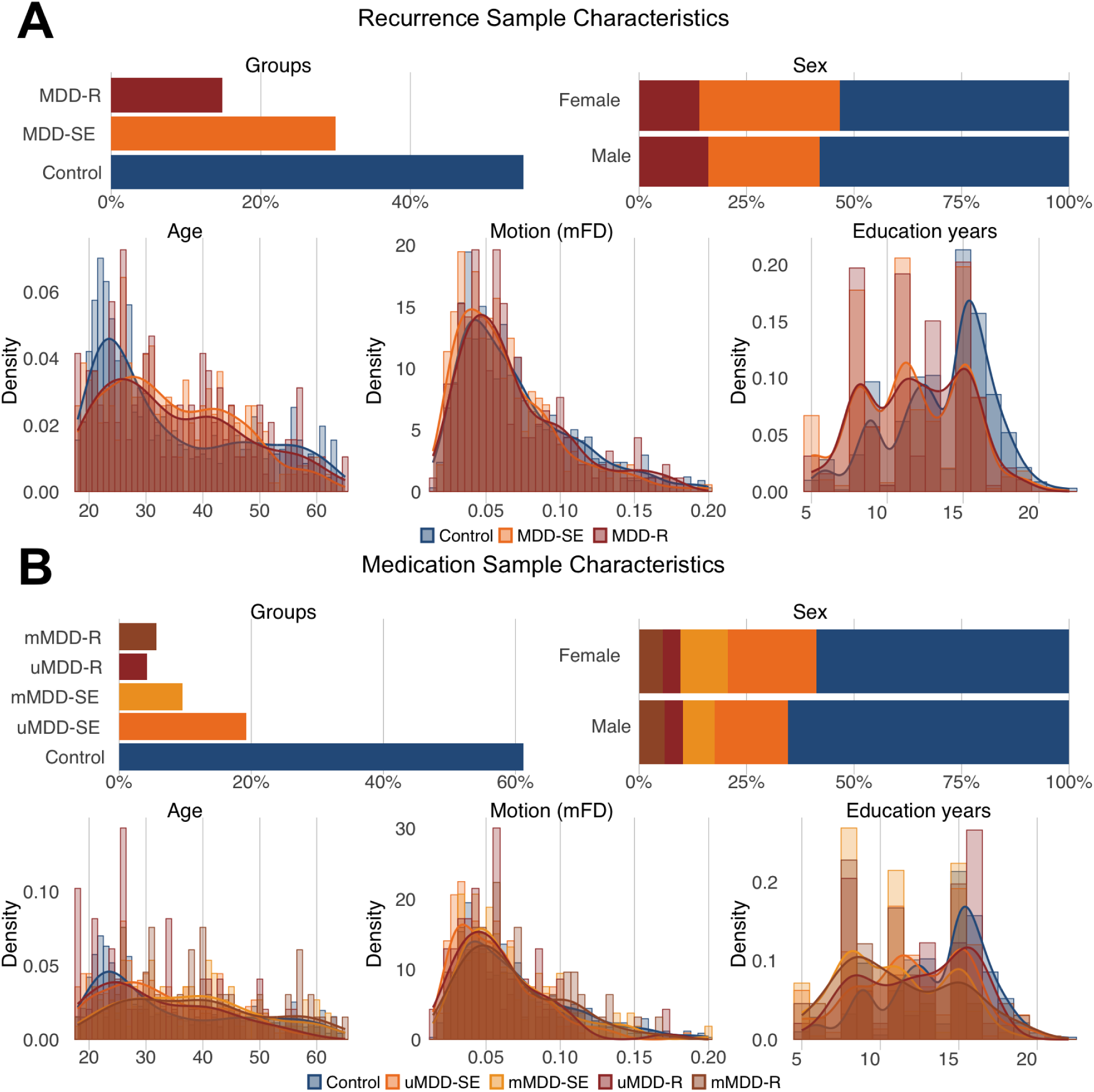
Demographic characteristics across the two main analytic subsamples. **(A)** Demographic variables for the recurrence-defined subsample (n = 1,296), **(B)** demographic variables for the medication-defined subsample (n = 1,166). Each panel includes group composition, age distribution, sex proportions, education years, and mean head motion (mFD). While small differences emerged across subsamples, all demographic variables were comparable between samples and included as covariates in the statistical models to control for potential confounding effects.

**Supplementary Fig. S3.**
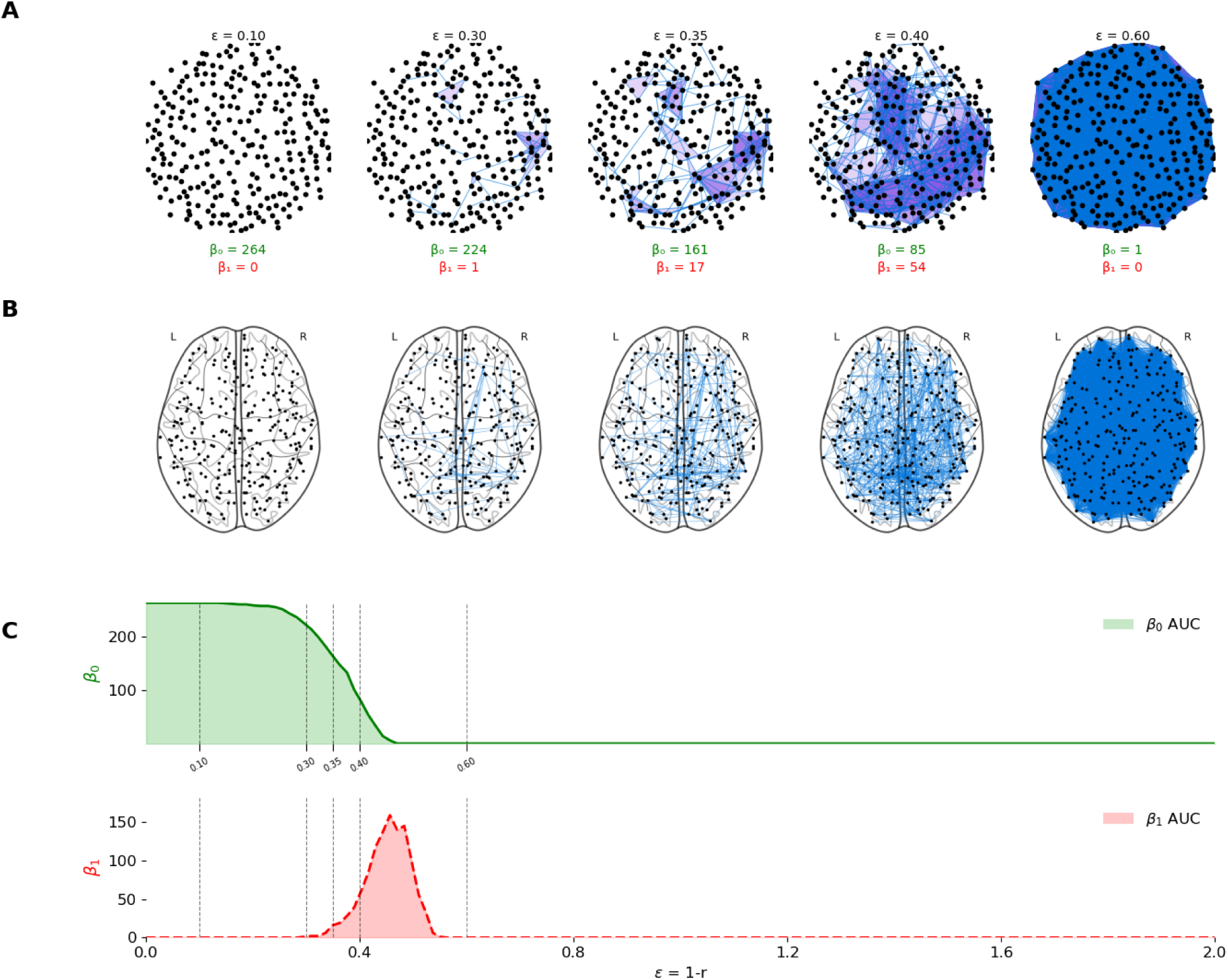
Topological filtration of a real functional connectome. **(A)** Visualization of the Vietoris–Rips filtration applied to a single subject. For solely visualization purposes, nodes are embedded in a two-dimensional space using multidimensional scaling (MDS) applied to the pairwise distance matrix. Edges (1-simplices) and filled triangles (2-simplices) are constructed based on the original distance matrix, with connections added whenever pairwise distances are less than or equal to the filtration value (ε). The resulting simplicial complexes illustrate the emergence and filling of topological features across scales. Betti numbers (β₀, β₁) computed at each ε indicate the number of connected components and hole-defining cycles at each filtration, respectively. **(B)** Corresponding anatomical connectivity graphs projected onto the Power-264 coordinates. At each ε, edges are added according to the same distance threshold, showing the evolving functional connectome in an anatomical space. **(C)** Betti curves as a function of the filtration parameter ε. The green curve (β₀) represents the number of connected components, while the red curve (β₁) represents the number of cycles. Dashed vertical lines indicate the filtration values shown in Panels **A–B**. Shaded regions correspond to the area under each curve (AUC), used as global summary measures of network connectivity (β₀ AUC) and higher-order topological organization (β₁ AUC).

**Supplementary Fig. S4.**
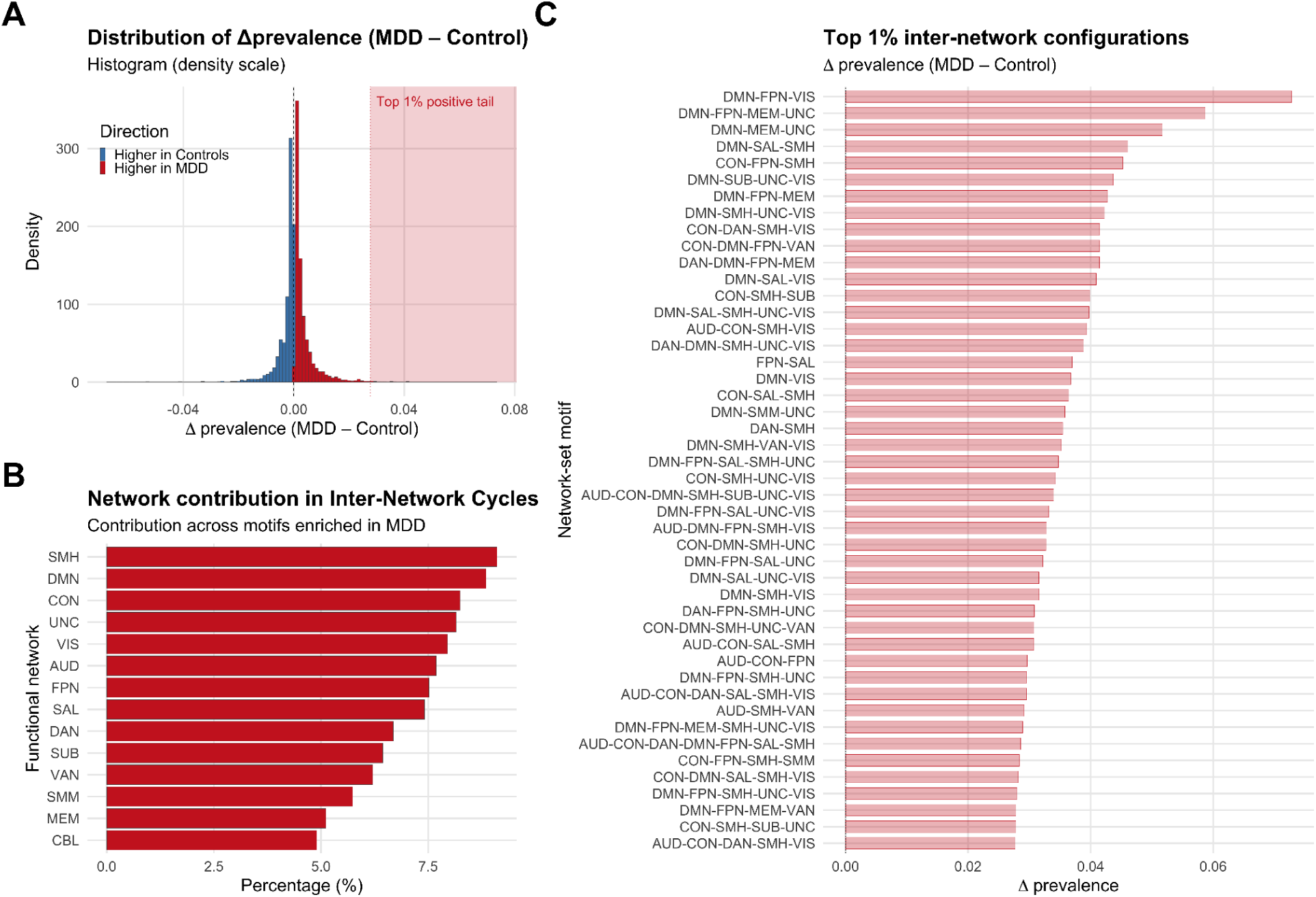
Network-set motifs of inter-network Betti-1 cycles filtering the 50% of most persistent cycles per subject. **(A)** Distribution of between-group differences in motif prevalence (Δprevalence = MDD – Control) across all inter-network motifs. The extreme positive tail (top 1%) is highlighted in red. **(B)** Decomposition of all motifs with higher prevalence in MDD (Δprevalence > 0) into their constituent single-networks, showing the total contribution of each functional system. Bars indicate the percentage of these motifs that include each canonical Power-264 network. The most represented networks were somatomotor-hand (SMH), default mode (DMN), cingulo-opercular (CON), uncertain/heteromodal (UNC), and visual (VIS), indicating that the network-involvement profile remained largely unchanged after restricting analyses to the 50% most persistent cycles. **(C)** Motifs in the extreme positive tail (top 1% of Δprevalence values), shown as illustrative examples of strongly MDD-biased inter-network configurations. This panel is descriptive and not inferential; motif identities shown here correspond to analyses restricted to the 50% most persistent cycles per subject. Network abbreviations (Power-264 atlas): SMH = Somatomotor-Hand; SMM = Somatomotor-Mouth; CON = Cingulo-opercular; DMN = Default Mode; FPN = Fronto-parietal; SAL = Salience; DAN = Dorsal Attention; VAN = Ventral Attention; VIS = Visual; AUD = Auditory; MEM = Memory Retrieval; SUB = Subcortical; CBL = Cerebellar; UNC = Uncertain.

